# Ephrin-A5 potentiates netrin-1 axon guidance by enhancing Neogenin availability

**DOI:** 10.1101/547299

**Authors:** L.-P. Croteau, T.-J. Kao, A. Kania

**Affiliations:** Institut de recherches cliniques de Montréal (IRCM), Montréal, QC, H2W 1R7, Canada; Department of Anatomy and Cell Biology and Division of Experimental Medicine, McGill University, Montréal, QC, H3A 2B2, Canada; Graduate Institute of Neural Regenerative Medicine, College of Medical Science and Technology, Taipei Medical University, Taipei 110, Taiwan; Technology and Center for Neurotrauma and Neuroregeneration, Taipei Medical University, Taipei 110, Taiwan

## Abstract

Axonal growth cones are guided by molecular cues in the extracellular environment. The mechanisms of combinatorial integration of guidance signals at the growth cone cell membrane are still being unravelled. Limb-innervating axons of vertebrate spinal lateral motor column (LMC) neurons are attracted to netrin-1 via its receptor, Neogenin, and are repelled from ephrin-A5 through its receptor EphA4. The presence of both cues elicits synergistic guidance of LMC axons, but the mechanism of this effect remains unknown. Using fluorescence immunohistochemistry, we show that ephrin-A5 increases LMC growth cone Neogenin protein levels and netrin-1 binding. This effect is enhanced by overexpressing EphA4 and is inhibited by blocking ephrin-A5-EphA4 binding. These effects have a functional consequence on LMC growth cone responses since bath addition of ephrin-A5 increases the responsiveness of LMC axons to netrin-1. Surprisingly, the overexpression of EphA4 lacking its cytoplasmic tail, also enhances Neogenin levels at the growth cone and potentiates LMC axon preference for growth on netrin-1. Since netrins and ephrins participate in a wide variety of biological processes, the enhancement of netrin-1 signalling by ephrins may have broad implications.

## Introduction

During nervous system assembly, neuronal wiring is specified by a rather limited set of axon guidance cues deployed at axonal trajectory decision points^1^. To achieve the high degree of complexity of neuronal connections found in even the simplest neural circuits, guidance signals act on axonal growth cones in a combinatorial manner, and are often integrated in a non-additive fashion such that their combined effects are different from their individual actions^2,3^. Unravelling the mechanisms underlying this interplay is an important question standing in the way of our complete understanding of how neuronal connections form.

One simple and well-described axon guidance decision is the selection of dorsal or ventral limb trajectory the base of the limb, executed by motor axons originating in the lateral motor column (LMC) of the spinal cord. The extension of the axons of the lateral LMC into the dorsal limb, and axons of the medial LMC into the ventral limb^4,5^ is specified by a number of axon guidance cues, including members of the ephrin and netrin protein families^6,7^. Ephrin ligands expressed in the limb repel LMC axons through their cognate receptors expressed in LMC axons: medial LMC axons express EphB tyrosine kinase receptors and are repelled from ephrin-Bs expressed in the dorsal limb, whereas lateral LMC axons express EphA receptors, including EphA4, and are repelled from ephrin-As, including ephrin-A5, in the ventral limb^8-11^. Reverse signalling from EphAs to ephrin-As, and its synergistic interaction with the GDNF and cRet/ GFRα1 ligand-receptor system are also important for lateral LMC axon guidance, revealing that even a simple binary axon guidance decision is subject to complex axon guidance cue interplay^12,13^.

Recent genetic and *in vitro* evidence argues that LMC axons entering the limb respond to netrin-1, a prototypical axon guidance cue that elicits axon attraction through its transmembrane receptors DCC and Neogenin and axon repulsion, through members of the UNC5 family^14-17^. Netrin-1 is expressed in the dorsal limb mesenchyme and lateral LMC axons show preference for growth over a netrin-1 containing substrate via the attractive netrin-1 receptor Neogenin. Medial LMC neurons avoid netrin-1 through the expression of Unc5c. Furthermore, netrin-1 acts synergistically with ephrins in LMC axon guidance such that medial LMC axons integrate netrin-1 and ephrin-B2 via a molecular complex containing Unc5c and EphB2. Its binding of netrin-1 and ephrin-B2 results in the increased activation of Src family of kinase effectors of netrin-1 and ephrin signalling, beyond that evoked by the presence of netrin-1 or ephrin-B2 alone. Lateral LMC axons also respond to netrin-1 and ephrin-A5 in a synergistic fashion, but the cellular and molecular mechanism of this effect remains elusive^18^.

Netrin-1 signalling specifies a wide variety of axon guidance decisions, often by acting in concert with other guidance cues^19,20^. For example, at the developing spinal cord midline, acting through their Robo receptors, Slit proteins silence netrin-1 attraction in commissural and motor axons^21,22^. In thalamocortical (TC) growth cones, Slit1 signalling via Robo1 and FLRT3 raises the levels of DCC allowing netrin-1 attraction, such that in the absence of Slit1, TC axons are unresponsive towards netrin-1^23, 24^. These and above studies suggest that netrin-1 signalling through its attractive receptors depends on the action of other axon guidance signals, prompting us to examine the mechanism of netrin-1 and ephrin-A integration by LMC axons. Our results demonstrate that ephrin-A5 induces an increase in Neogenin abundance in LMC growth cones through its receptor EphA4, sensitizing lateral LMC axons to netrin-1. This effect occurs in the absence of the intracellular signalling tail of EphA4, demonstrating that lateral LMC axon repulsion from ephrin-A5 and sensitization to netrin-1 occur through molecularly distinct pathways.

## Results

### Ephrin-A5 sensitizes lateral LMC axons to netrin-1

At the time of their growth into the limb mesenchyme, chick lateral LMC axons respond synergistically to the presence of ephrin-A5 and netrin-1: while these axons are insensitive to low concentrations of either netrin-1 or ephrin-A5, when challenged simultaneously with stripes containing low concentrations of netrin-1 and ephrin-A5, lateral LMC axons exhibit a robust growth on netrin-1 stripes^18^. We envisaged two possible mechanisms explaining this behaviour: (1) netrin-1 sensitizes lateral LMC axons to ephrin-A5 avoidance or (2) ephrin-A5 sensitizes lateral LMC axons to netrin-1 attraction. To distinguish between these, we studied the *in vitro* behaviour of LMC neurons explanted from chick spinal cords within the developmental window in which LMC axons innervate the limbs (Hamburger-Hamilton stage (HH st.) 25-26^25^). Such LMC explants were placed on carpets of two alternating stripes containing (1) a mixture of Cy3 secondary antibody and either ephrin-A5-Fc (referred as ephrin-A5 subsequently) or netrin-1 and (2) stripes containing Fc protein, with and without bath application of netrin-1 or ephrin-A5 (**Figure 1)**. Lateral LMC axons were visualized via EphA4 expression, a lateral LMC marker^26^, and their stripe preference outgrowth was scored as previously^27^. When lateral LMC axons were challenged with alternating ephrin-A5 and Fc stripes (ephrin-A5 / Fc) with or without low concentration of bath-applied netrin-1 (10 ng/mL), no avoidance of ephrin-A5 stripes was observed (**Figure 1c, d**; 52% and 50% respectively, p=0.57). In contrast, while lateral LMC axons challenged with alternating netrin-1 (10 ng/mL) and Fc stripes (netrin-1/Fc) showed no growth preference (**Figure 1a**, 47% on netrin-1, p>0.05), in the presence of bath-applied ephrin-A5 (50 ng/mL), lateral LMC axons robustly preferred netrin-1 over Fc stripes (**Figure 1b**, netrin-1 67% on netrin-1, p<0.001). These results suggest that ephrin-A5 sensitizes LMC axons to netrin-1.

**Figure 1.**
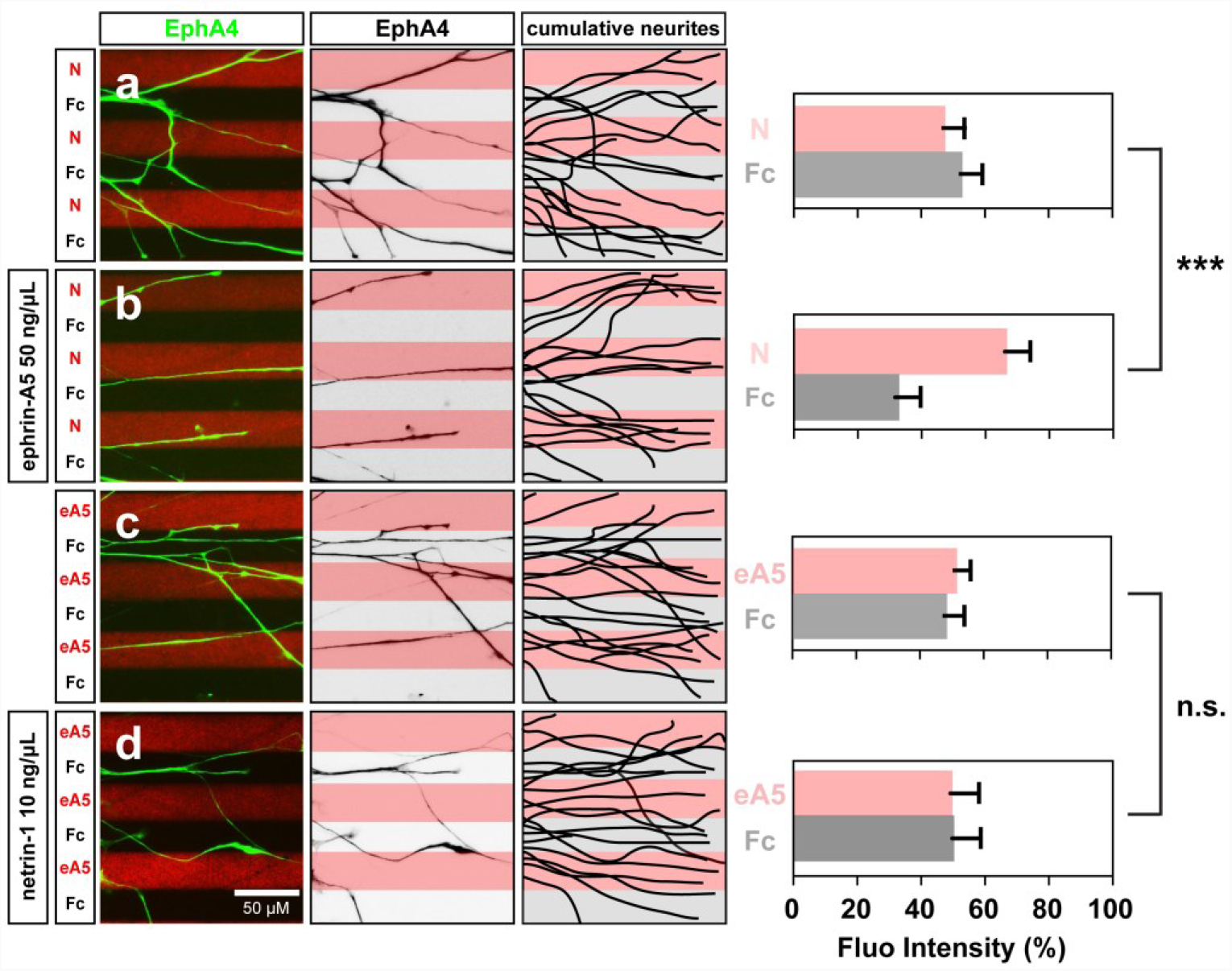
Lateral LMC axon growth responses to netrin-1 are sensitised by ephrin-A5. Axon outgrowth preference on protein stripes exhibited by lateral LMC axons. left panels: explanted lateral (EphA4+) LMC neurites on netrin-1 (N)/Fc (a,b) or ephrin-A5 (eA5)/Fc stripes (c,d) with or without bath treatment of ephrin-A5 (b) or netrin-1 (d). Middle panels: inverted images of EphA4 signals shown at left panels. Right panels: superimposed images of five explants from each experimental group representing the distribution of lateral LMC neurites. Quantification of lateral LMC neurites on first (pink) and second (pale) stripes expressed as a percentage of total EphA4 signals. Noted that no preference is detected when lateral LMC neurites are challenged with stripes of low levels of netrin-1 (10 ng/mL) or ephrin-A5 (50 ng/mL). Minimal number of neurites: 81. Minimal number of explants: 12. N, netrin-1; eA5: ephrin-A5; error bars = SD; *** = P<0.001; statistical significance computed using Mann-Whitney U test.

### Ephrin-A5 increases Neogenin and EphA4 protein levels in LMC growth cones

Chicken motor axon attraction to netrin-1 is mediated through its attractive receptor Neogenin^18^. To investigate the possibility that ephrin-A5 sensitises LMC axons to netrin-1 by increasing Neogenin abundance in growth cones, we measured the relative Neogenin protein levels in LMC growth cones by immunofluorescence (IF) using an anti-Neogenin affinity-purified polyclonal antiserum directed against the extracellular domain of Neogenin (anti-Neogenin polyclonal antibody)^18^. HH st. 24-25 LMC explants were incubated over-night and treated for 20’ with various concentrations of ephrin-A5 (**Figure 2a-e**). The average area of the growth cones selected for analysis did not differ between treatments (**Supplemental figure 1**). However, compared to Fc control treatment, application of pre-clustered ephrin-A5 at 50 and 100 ng/mL, resulted in, respectively, 1.7-fold and 2-fold increases in the levels of Neogenin IF (**Figure 2a**; p= 0.03, and p=0.01), while higher ephrin-A5 concentration did not cause a significant rise in Neogenin IF (**Figure 2a**; 250 ng/ml: p=0.08, 500 ng/mL: p=0.93). Similar Neogenin IF increase was also detected using a polyclonal anti-Neogenin antiserum directed against the cytoplasmic tail and extracellular Neogenin: ephrin-A5 100 ng/mL exposure resulted in a 1.7-fold increase in Neogenin IF (**Figure 3a-q**; intracellular Neogenin vs. MN media: p=0.01; vs. extracellular Neogenin: p=0.16).

**Figure 2.**
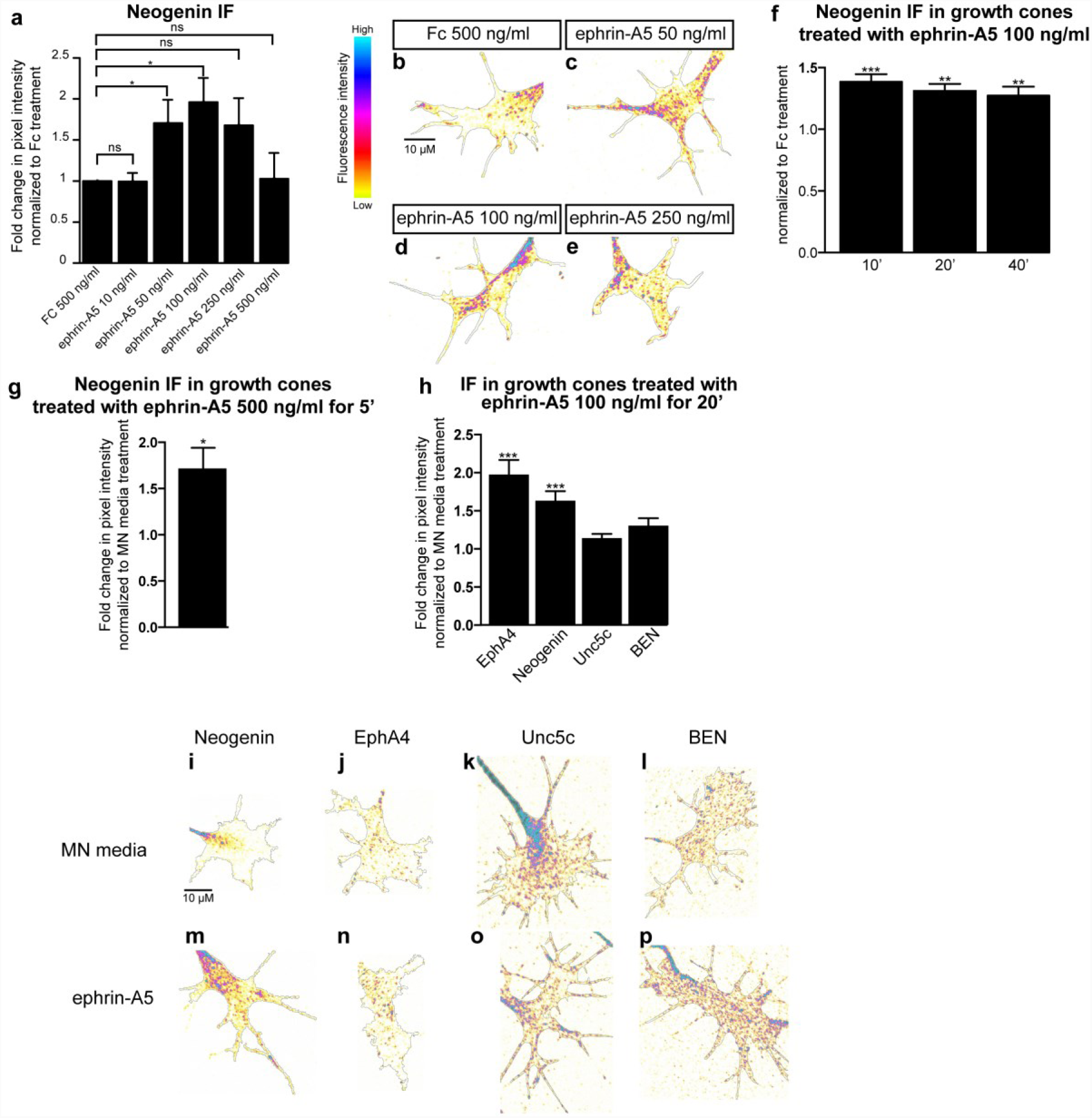
Ephrin-A5 increases Neogenin and EphA4 protein levels in LMC growth cones. Mean Neogenin IF in LMC explants treated 15’ with either a control solution containing Fc at 500 ng/mL or ephrin-A5 at concentrations ranging from 10 to 500 ng/mL. Ephrin-A5 at 50 and 100 ng/mL results in increased levels of Neogenin IF (ephrin-A5 50 ng/mL: 1.71-fold, p= 0.038; ephrin-A5 100ng/mL: 1.96-fold, p=0.011). (b-e) Examples of Neogenin IF in growth cones quantified in (a). (f) Mean Neogenin IF in growth cones of LMC explants treated with either Fc or ephrin-A5 at 100 ng/mL for 10’ 20’ and 40’. A 10’ exposure to ephrin-A5 is sufficient to increase Neogenin IF (1.39-fold, p= 0.0007) and the increase is maintained after 20’and 40’ (1.31-fold, p= 0.0017; 1.27-fold, p= 0.0095 respectively). (g) A 5’ exposure to ephrin-A5 at 100 ng/mL is sufficient for increasing the mean Neogenin IF in LMC growth cones (1.71-fold, p=0.0228). (h) Quantification of mean Neogenin, EphA4, Unc5c and BEN IF levels in growth cones exposed to either MN media or ephrin-A5 at 100 ng/mL for 20’. Ephrin-A5 induced an increase in Neogenin and EphA4 IF (1.64-fold, p < 0.0001; 1.90-fold,p= 0.0001 respectively), Unc5c and BEN IF levels did not significantly differ (p=0.0806 and p= 0.1391 respectively). (i,p) Examples of growth cones quantified in (h). Data are shown as mean ±SEM, statistical significance was tested using a two-tailed unpaired sample t-test. (a) Fc 500 ng/mL N=5, ephrin-A5 10 ng/mL N=3, eprhrin-A5 50 ng/mL N=5, ephrin-A5 100 ng/mL N=5, ephrin-A5 250 ng/mL N=5, ephrin-A5 500 ng/mL N=3; (f) Fc 100 ng/mL 40’ N=4,ephrin-A5 100 ng/mL 10’ N=4, ephrin-A5 100 ng/mL 20’ N=4, ephrin-A5 100 ng/mL 40’ N=4; (g) Fc 500 ng/mL 5’ N=4, ephrin-A5 500 ng/mL 5’ N=4; n: EphA4 N=16,, Neogenin N=13, BEN N=5, Unc5c N=10. Values for the total number of growth cones and SEM values for each treatment for this figure and subsequent figures are provided in the supplementary Excel file.

**Figure 3.**
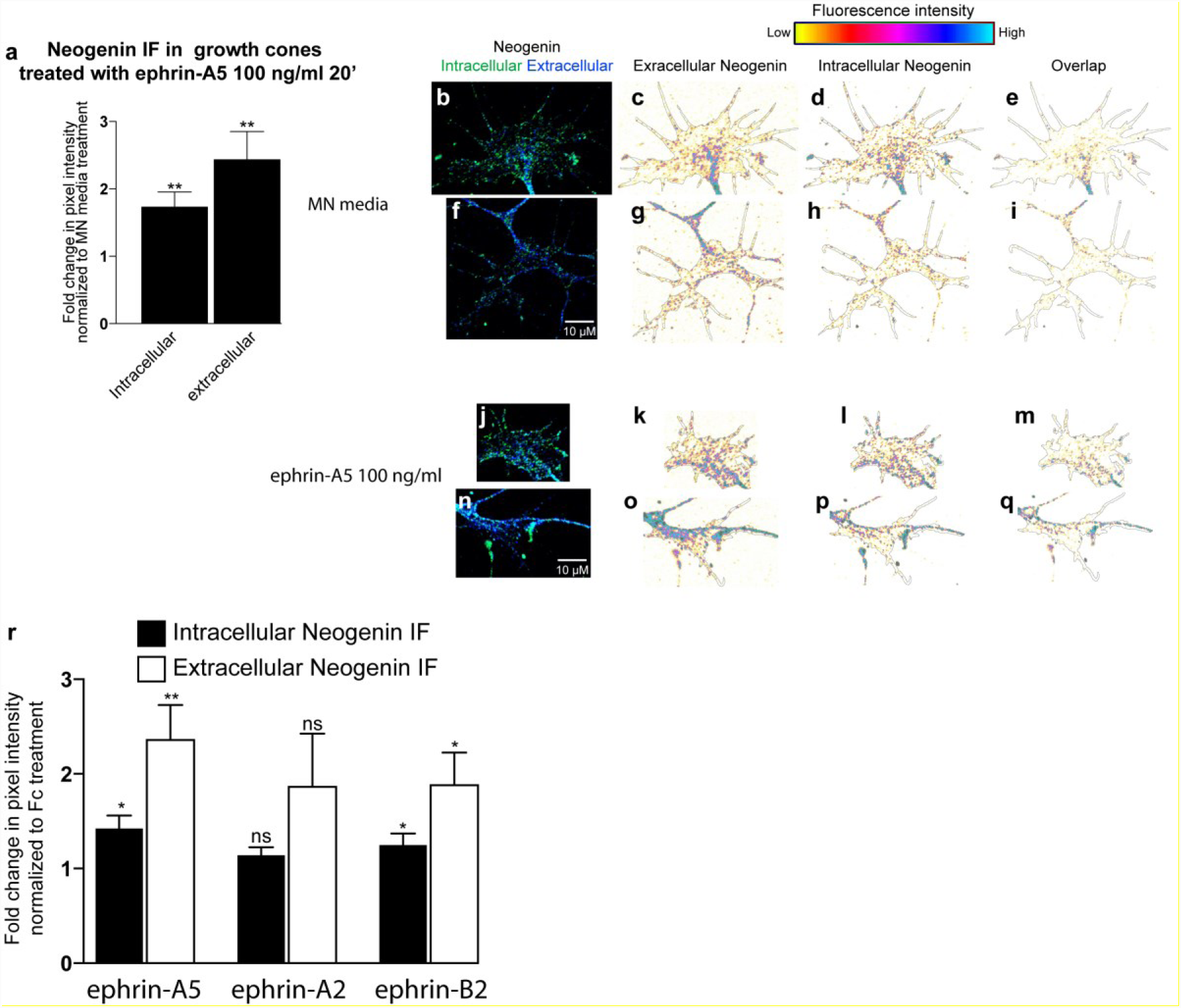
Ephrin-A5 and Ephrin-B2 increase both intracellular and extracellular Neogenin IF. (a-q) Compared to MN media, ephrin-A5 100 ng/mL increases both intracellular and extracellular Neogenin IF in LMC growth cones (1.74-fold, p=0,005; 2.44-fold, p=0,004 respectively, N=7). (b-q) Examples of growth cones quantified in (a). (r) A 20’ exposure to ephrin-A5, ephrin-B2 but not ephrin-A2 at 100 ng/mL increases intra. and extra. Neogenin IF in LMC growth cones (ephrin-A5 intra. Neogenin IF: 1.42-fold, p=0,006; ephrin-A5 extra. Neogenin IF: 2.36-fold, p=0.002; ephrin-A2 intra. Neogenin IF: 1.14-fold, p=0.064; ephrin-A2 extra. Neogenin IF: 1.86-fold, p=0.078; ephrin-B2 intra. Neogenin IF: 1.25-fold, p=0.034; ephrin-B2 extra. Neogenin IF: 1.88-fold, p=0.015. N=6. Data are shown as mean ±SEM, statistical significance was tested using a two-tailed (a) and one-tailed (r,s) unpaired sample t-test.

To characterize the dynamics of the ephrin-A5-induced rise in Neogenin IF, explants were subject to 100 ng/mL ephrin-A5 for 10’, 20’ and 40’. The Neogenin IF increase occurred within 10’ of exposure to ephrin-A5 and remained as such after 40’ (**Figure 2f**). Furthermore, even a 5’ exposure to ephrin-A5 at 500 ng/mL resulted in a 1.71-fold increase in Neogenin IF (**Figure 2g;** p=0.02).

To assess the specificity of these effects, we also quantified the consequence of ephrin-A5 treatment of LMC growth cones on IF levels of the ephrin-A5 receptor EphA4, the netrin-1 repulsive receptor Unc5c, as well as the surface glycoprotein BEN^10,18,28^. Ephrin-A5 addition significantly increased EphA4 IF, possibly be due to an increase in EphA4 trafficking to the growth cone plasma membrane resulting from ephrin-A5-EphA4 interactions^29^. It is also possible that the oligomerization of EphA4 when bound to clusters of ephrin-A5 could contribute to increasing EphA4 IF^30^ (1.9-fold, p=0.0001) without affecting Unc5c or BEN IF levels (**Figure 2h**). Ephrin-A5 treatment did not significantly alter the area of growth cones included in the IF analysis (**Supplementary Figure 1a-d**). Together, these data suggest that the increased LMC growth cone sensitivity to netrin-1 following ephrin-A5 treatment could be due to an increase in Neogenin abundance.

### Ephrin-B2 increases Neogenin protein levels in LMC growth cones

Ephrin-A2 and ephrin-B2 also guide LMC axons^11,12, 18^, prompting us to determine whether these two ephrins can also influence Neogenin levels in LMC growth cones. As above, explanted LMC growth cones were treated for 20’ with either Fc, ephrin-A5, ephrin-A2 or ephrin-B2 at 100 ng/mL, followed by a quantification of Neogenin IF (**Figure 3r,s**). Both, ephrin-A5 and ephrin-B2 but not ephrin-A2 caused an increase in Neogenin IF (**Figure 3r,s**; ephrin-A5 intra.: 1.4-fold, p=0.006, A5 extra.: 2.3-fold, p=0.002; ephrin-B2 intra.: 1.25-fold, p=0.03 B2 extra.: 1.88-fold, p=0.015). These results suggest that ephrins differ in their ability to increase Neogenin IF in LMC growth cones.

### Ephrin-A5 increases Neogenin protein abundance on the surface of growth cones

Ephrin-A5 induced LMC axon sensitization to netrin-1 may result from increased abundance of Neogenin on the surface of growth cones. To test this possibility, we applied the anti-Neogenin polyclonal antiserum with either ephrin-A5 at 50 ng/mL or MN media as control, to live LMC growth cones for 20’, followed by standard fixation. Application of an antibody against the intracellular protein β III tubulin, did not result in any labelling, suggesting that this treatment results in specific detection of cell surface epitopes (**Supplementary Figure 2a-h**). A 20’ treatment with ephrin-A5 resulted in a 1.6-fold increase in surface Neogenin IF (**Figure 4a-c**; p=0.04), arguing that ephrin-A5 application results in increased of Neogenin protein levels on the surface of LMC growth cones.

**Figure 4.**
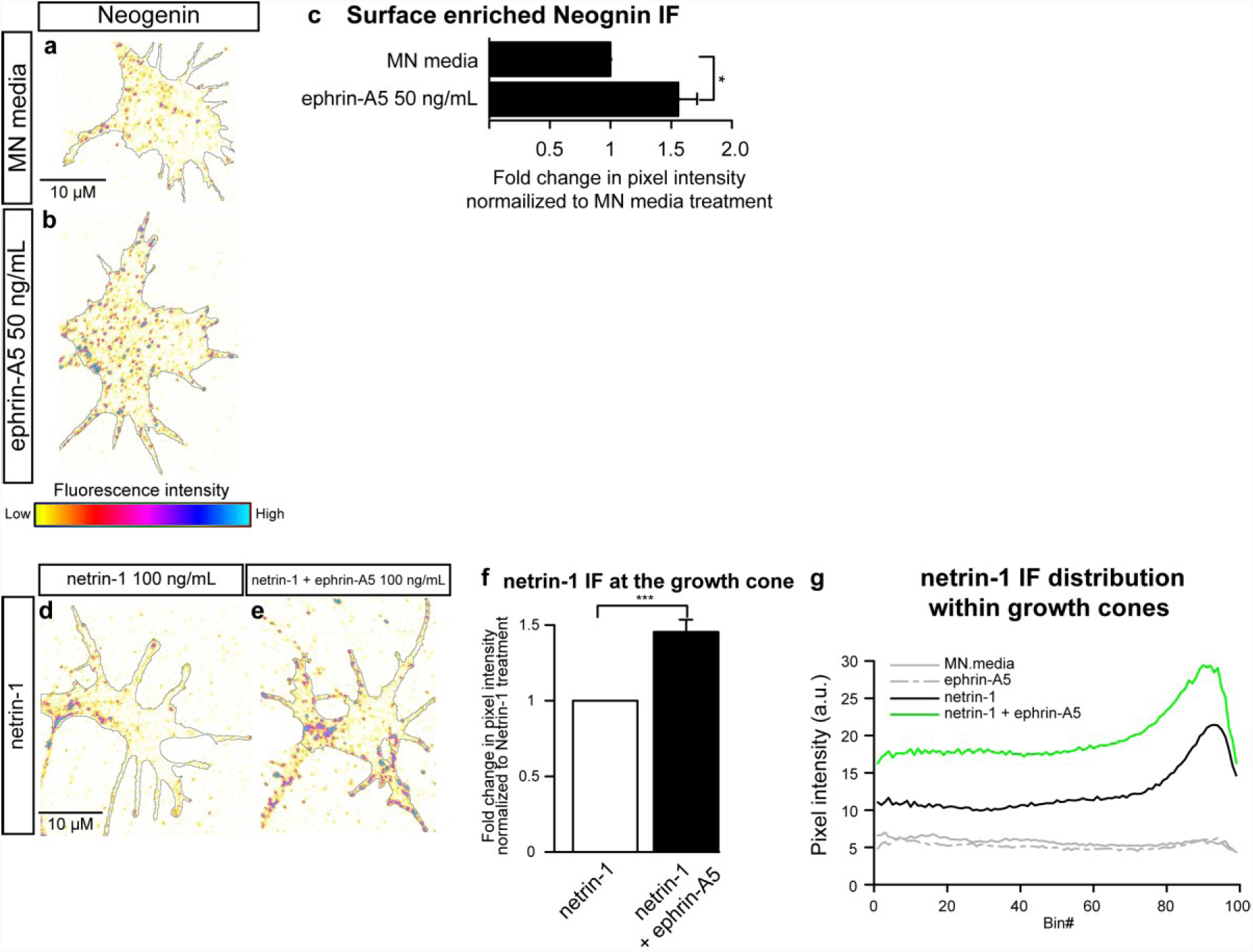
Exposure to ephrin-A5 increases surface enriched Neogenin protein levels and netrin-1 binding to LMC growth cones. Examples of surface-enriched Neogenin in MN media (a) or ephrin-A5 50 ng/mL (b) treated LMC growth cones. (c) Quantification of surface enriched Neogenin IF in LMC growth cones treated with either MN media or ephrin-A5 50 ng/mL for 20’, ephrin-A5 increases surface enriched Neogenin IF 1.56-fold, p=0.037. (d,e) Examples of netrin-1 IF in LMC growth cones treated with either netrin-1 (d) or a combination of netrin-1 and ephrin-A5 (100 ng/mL). (f) Quantification of netrin-1 IF in growth cones treated with netrin-1 or netrin-1 + ephrin-A5, addition of ephrin-A5 results in a 1.45-fold increase in netrin-1 IF (p=0.0003). (d) Graph depicting the distribution of netrin-1 IF in growth cones of explants treated with either MN media, ephrin-A5, netrin-1 or netrin-1 + ephrin-A5. Bin #1 coincides with the geometric centre of the growth cone and Bin# 99 coincides with the perimeter of the growth cone. Data are shown as mean ±SEM, statistical significance was tested using a two-tailed unpaired sample t-test. (c) N=4 (f,g) MN media: N=3; netrin-1: N=6; ephrin-A5: N=6; netrin-1 + ephrin-A5: N=6.

### Ephrin-A5 enhances netrin-1 binding in growth cones

We next sought to determine whether increased LMC growth cone surface Neogenin IF levels might result in increased netrin-1 binding to LMC growth cones. LMC explants were incubated with netrin-1 and ephrin-A5 or netrin-1 alone as control. After fixation, a monoclonal antibody against netrin-1 was used to detect relative netrin-1 binding (**Figure 4d-g**). Compared to a netrin-1 treatment, the addition of ephrin-A5 resulted in increased levels of netrin-1 IF in LMC growth cones (**Figure 4d-g**, 1.4-fold increase, p= 0.0003), without a change in growth cone size (**Supplementary Figure 1**). These result suggest that the sensitization of LMCl axons to netrin-1 by ephrin-A5 (**Figure 1a,b**) may be a consequence of enhanced netrin-1 binding in LMC growth cones.

### Ephrin-A5 does not change Neogenin abundance in growth cones of dorsal spinal cord neurons

To test the susceptibility of other neuronal cell types to an ephrin-A5 induced increase in Neogenin IF, HH st. 24-25 dorsal lumbar spinal cord explants, which express Neogenin^31,32^, were incubated overnight and subjected to a 20’ treatment with either MN media or ephrin-A5 100 ng/mL followed by an EphA4 and Neogenin IF analysis. Neither EphA4 or Neogenin IF was altered by ephrin-A5 (**Figure 5a**; p=0.5, p=0.3 respectively). To compare the relative levels of EphA4 and Neogenin between LMC and dorsal spinal cord growth cones, LMC and dorsal explants from the same spinal cord segments were cultured and analyzed as described above (**Figure 5b-k**). The levels of both EphA4 and Neogenin IF were higher in MN media-treated dorsal explants relative to LMC explants (**Figure 5b**; EphA4: 3.1-fold, p<0.0001, **Figure 5c**; Neogenin: 4.1-fold, p=0.005). These data suggest that the ephrin-A5 elevation of Neogenin levels occurs only in select population of neuronal growth cones (**Figure 5b,c**).

**Figure 5.**
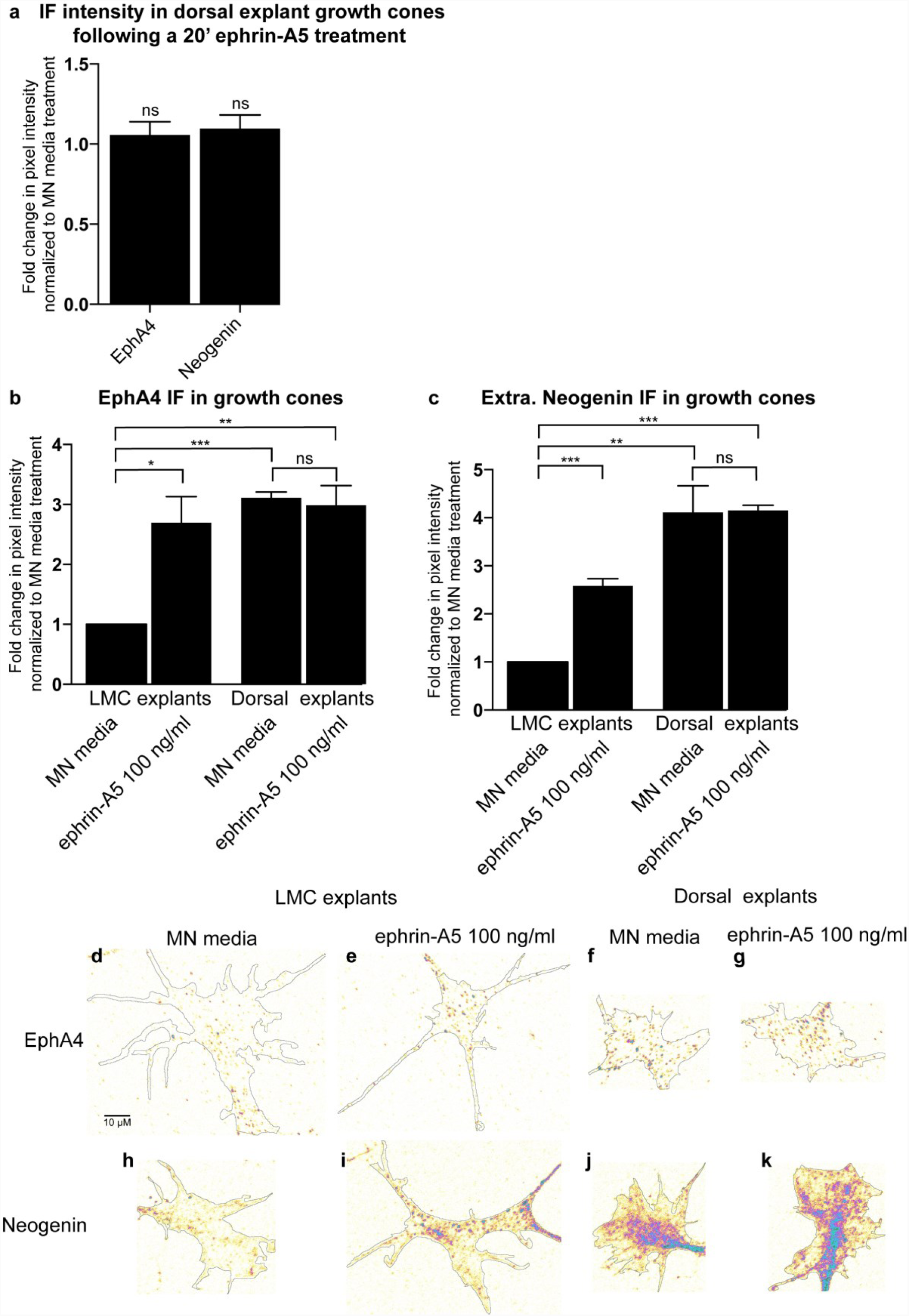
Ephrin-A5 fails to increase Neogenin or EphA4 protein levels in growth cones of dorsal spinal cord explants. (a) Quantification of the mean Neogenin and Epha4 IF in growth cones of dorsal lumbar spinal cord explants treated with ephrin-A5 100 ng/mL for 20’ (p=0.835 and p=0.825 respectively). (b,c) A comparison of Neogenin and EphA4 IF levels between growth cones of LMC and dorsal spinal cord explants shows that EphA4 and Neogenin IF is greater in dorsal explants (3.11-fold, p= 0.00003 and 4.11-fold, p=0.0049 respectively). Whereas ephrin-A5 increases EphA4 and Neogenin IF levels in LMC explants (2.69-fold, p=0,018; 2.58-fold, p=0,0005 respectively), EphA4 and Neogenin IF in dorsal explants remains unchanged (p=0.734 and p= 0.949). (d,K) Examples of growth cones quantified in (b,c). Data are shown as mean ±SEM, statistical significance was tested using a two-tailed unpaired sample t-test. (a) MN media N=6, Ephrin-A5 N=6; (b,c) MN explants / MN media N=3, MN explants / Ephrin-A5 N=3, Dorsal explants / MN media N=3, Dorsal explants / Ephrin-A5 N=3.

### Ephrin-A5 induces an increase in the co-localization of Neogenin and EphA4 IF in LMC growth cones

Following ephrin-A5 treatments, we noticed the formation of dense EphA4 IF puncta in LMC growth cones (**Figure 2n**), likely resulting from ligand-induced EphA4 clustering^30^. To investigate the possibility that the ephrin-A5 increase in EphA4 and Neogenin IF may be spatially correlated, we analysed the co-localization of Neogenin and EphA4 IF in LMC growth cones. LMC neuron explants were treated with either MN media, ephrin-A5 or netrin-1 (at 100 ng/mL) followed by immunostaining for EphA4 and either Neogenin, BEN or Unc5C in LMC growth cones. Treatment with ephrin-A5 resulted in an increase in the co-localisation of Neogenin with EphA4 IF, but not with Unc5c or BEN IF (**Figure 6a**, 1.73-fold, p=0.0004; 1.55-fold, p=0.04 respectively), suggesting that the increase in Neogenin protein levels may be occurring in close proximity to ephrin-A5-EphA4 clusters (**Figure 6b-i**).

**Figure 6.**
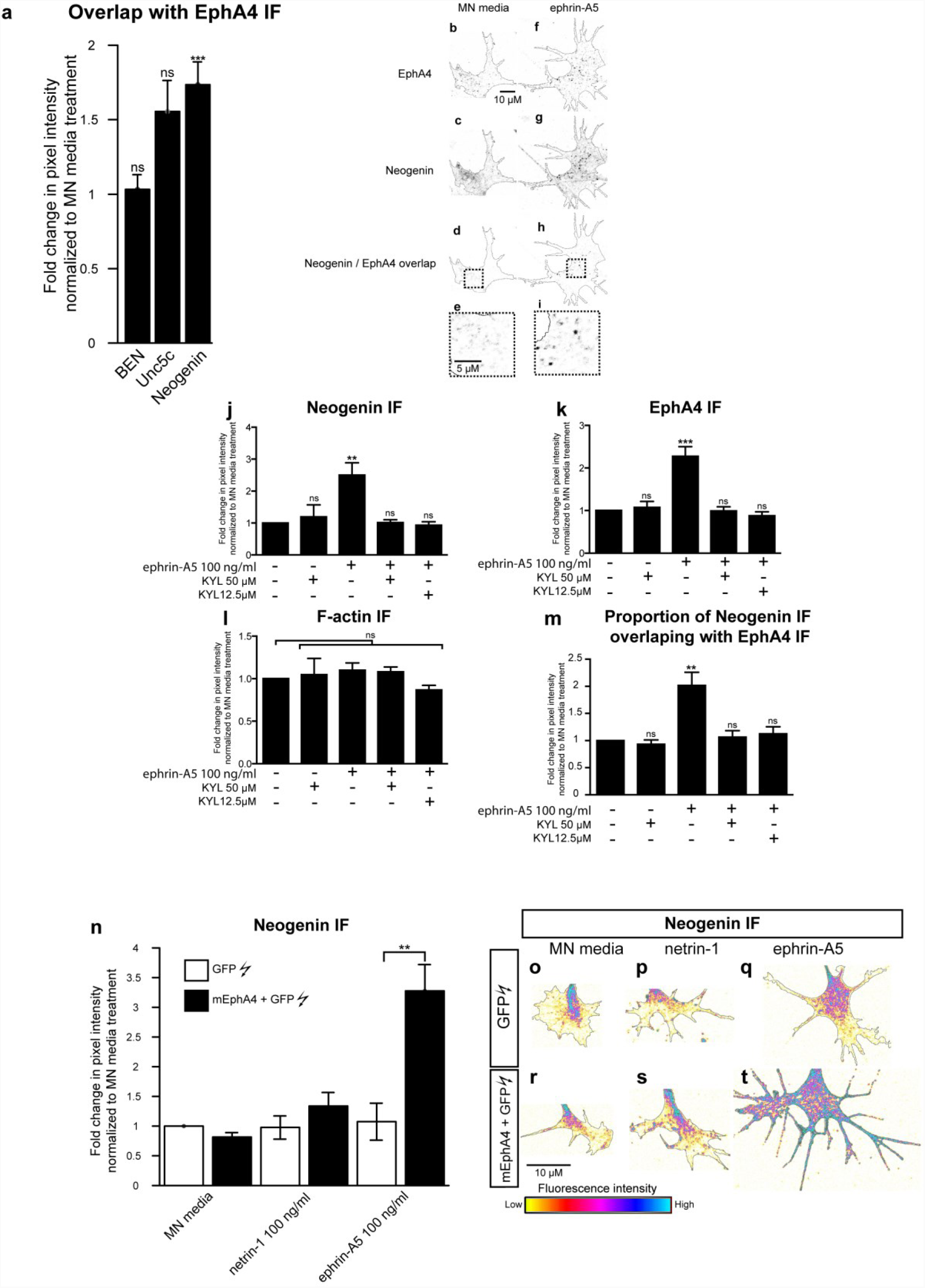
Involvement of EphA4 in the ephrin-A5-induced increase in Neogenin IF. Quantification of the proportion of Neogenin, Unc5c and BEN IF overlap with EphA4 IF in thresholded images of growth cones that were treated for 20’ with MN media ephrin-A5 at 100 ng/mL. Ephrin-A5 treatment resulted in a 1.73 - fold increase in the proportion of Neogenin/EphA4 IF overlap (p=0.0007) without significantly altering Unc5c/EphA4 or BEN/EphA4 IF overlap (p>0.05). (b,i) Examples of thresholded images of growth cones immunostained for EphA4 (b,f) and Neogenin (c, g) and treated with MN media (b-e) or ephrin-A5 (e-i). Panels (d) and (h) show the resulting overlap in Neogenin/EphA4 signal in MN media and ephrin-A5 treated explants respectively, panels (e,i) are magnified images of boxed regions in (d,h). (j-m) LMC explants were incubated with either MN media, KYL 12.5 µM or KYL 50 µM for 20’ prior to being treated with either MN media or ephrin-A5 at 100 ng/mL for 20’ followed by Neogenin, EphA4 and F-actin immunostaining. (j-l) Quantification of Neogenin (j), EphA4 (k) or F-actin (l) IF in growth cones normalized to MN media treatment. (m) Quantification of the proportion of Neogenin IF overlapping with EphA4 IF normalized to MN media treatment. (j) ** p=0.0034, (k) ***p=0.0003, (m) **p=0.0085. (n-t) LMC explants from chick spinal cords electroporated with either a GFP expression plasmid alone or in combination with a mEphA4 expression plasmid were subject to a 20’ treatment of either MN media, netrin-1 or ephrin-A5 at 100 ng/mL. (n) Quantification of Neogenin IF in growth cones shows that in explants treated with ephrin-A5 and overexpressing EphA4, Neogenin levels in growth cones are 3.28-fold higher than in explants expressing GFP alone (** p=0.0022). (o-t) Examples of Neogenin IF in growth cones quantified in (n). Data are shown as mean ±SEM, statistical significance was tested using a two-tailed unpaired sample t-test. The number of experiments and growth cones for each treatment are in the supplementary Excel file.

### Ephrin-A5 - EphA4 binding is required for the growth cone elevation of Neogenin by ephrin-A5

The ephrin-A5 induced increase in EphA4 IF and Neogenin-EphA4 IF co-localization in LMC growth cones suggests that the increase in Neogenin IF may be dependent on ephrin-A5-EphA4 interactions. To test this idea, ephrin-A5-EphA4 interactions were blocked using a 12-amino-acid peptide (KYL) that selectively binds EphA4 at its ephrin binding domain and inhibits ephrin-A5/EphA4 interactions without affecting ephrin-A5 binding to other Eph receptors^33^. LMC explants were treated with either MN media, KYL peptide at 50 µM or KYL peptide at 12.5 µM for 20’ prior to a 20’ exposure to either MN media or ephrin-A5 at 100 ng/mL (**Figure 6j-m**). The pre-incubation of LMC explants with either KYL at 12.5 or 50 µM abolished the ephrin-A5 induced increase in Neogenin IF, EphA4 IF and in the co-incidence of Neogenin and EphA4 IF (**Figure 6j-m**). These results argue that the LMC growth cone increase in Neogenin IF caused by ephrin-A5, requires its binding to EphA4.

### EphA4 potentiates the ephrin-A5-led increase in Neogenin abundance

Since blocking EphA4 function resulted in the attenuation of ephrin-A5-induced Neogenin IF increase, we reasoned that increasing EphA4 expression levels in LMC growth cones would increase the ephrin-A5-mediated induction of Neogenin. To test this idea, *GFP* expression plasmids alone or together with mouse *EphA4* expression plasmids were introduced into chick neural tubes at HH st. 18-19 by *in ovo* electroporation^10,34^. GFP-expressing LMC neurons were explanted at HH st. 24-25, treated with MN media, netrin-1 or ephrin-A5 (either at 100 mg/mL) and Neogenin IF levels were quantified in GFP+ growth cones (**Figure 6n-t**). Ephrin-A5 treatment of EphA4 and GFP overexpressing LMC neurons led to a 3-fold increase in Neogenin IF compared to LMC neurons overexpressing GFP only (**Figure 6n,q,t**, p=0.007), arguing that EphA4 is sufficient to potentiate the ephrin-A5-induced increase in Neogenin IF.

To gain further insights in the mechanism behind the ephrin-A5-dependent increase in Neogenin IF, we used pharmacological inhibitors of specific cellular processes, subjecting LMC explants overexpressing EphA4 to these for 20’ prior to ephrin-A5 and netrin-1 treatment, followed by growth cone Neogenin IF quantification (**Supplementary Figure 2i**). To assess the possible role of PKA, we used the PKA inhibitor KT5720 (5 µM,^35^), to block protein synthesis we used anisomycin (80 µM,^36^). Proteasomal and lysosomal Neogenin degradation was blocked by MG132 (20 µM,^37^) and chloroquine (20 µM,^38^), respectively. Since both EphA4 and Neogenin can be cleaved by γ-secretase^39,40^, LMC explants were treated with the γ-secretase inhibitor DAPT^41^. Finally, since Src family kinases are activated downstream of both EphA4 and Netrin-1 receptors^42-46^, LMC explants were treated with the Src family inhibitor SU6656^47^. None of these treatments resulted in a significant increase in Neogenin IF when treated with MN media or attenuated the increase in Neogenin levels following ephrin-A5 treatment (**Supplementary Figure2i**).

### The ephrin-A5-induced upregulation of Neogenin does not depend on protein synthesis or proteolysis

The above lack of inhibitor effects could be because these were used in the context of LMC growth cones overexpressing EphA4, whose elevated signalling might be impervious to a partial pharmacological blockade, prompting us to re-examine protein and protease inhibition in the context of wild type LMC explants. Thus, if either process is involved in the augmentation of Neogenin protein levels by ephrin-A5, their blockade may also affect Neogenin levels. LMC explants were pre-treated for 20’ with either DMSO or the protein synthesis inhibitor cycloheximide (CHX) 50 µM, followed by a 5’ treatment with either Fc or ephrin-A5 500 ng/mL (**Figure 7a**). The ephrin-A5-induced increase in Neogenin IF was evident even in the presence of CHX and did not significantly differ from DMSO pre-treatment (**Figure 7a**, p=0.418). To investigate the short-term effects of blocking proteolytic cleavage on Neogenin and EphA4 levels, LMC explants were subjected to a broad-spectrum commercial protease inhibitor cocktail^48^ and after a 20’ treatment, F-actin, EphA4 and Neogenin IF were not significantly altered relative to DMSO treatment (**Figure 7b**, 1.19-fold, p=0.08; 1.23-fold, p=0.06; 1.28-fold, p=0.08 respectively). The ectodomain cleavage of Dcc, a member of the immunoglobulin domain family closely related to Neogenin ^32^, can be blocked by the broad-spectrum metalloprotease inhibitor GM6001^49^. However, relative to DMSO, a 20’ treatment with GM6001 failed to increase EphA4 and Neogenin IF (**Supplementary Figure 2l**, 1.25-fold p=0.1; 1.46-fold, p= 0.07 respectively). Furthermore, since potential proprotein convertases cleavage sites are present in both EphA4 and Neogenin^50^, we treated LMC explants with ephrin-A5 and the broad spectrum proprotein convertase inhibitor RVKR^51^. This resulted in 1.31-fold increase in Neogenin IF without affecting EphA4 IF (**Supplementary Figure 2k**, p=0.01 and p=0.2 respectively). Altogether, these results suggest that a 20’ inhibition of protein synthesis or proteolytic cleavage does not substantially increase Neogenin abundance in LMC growth cones.

**Figure 7.**
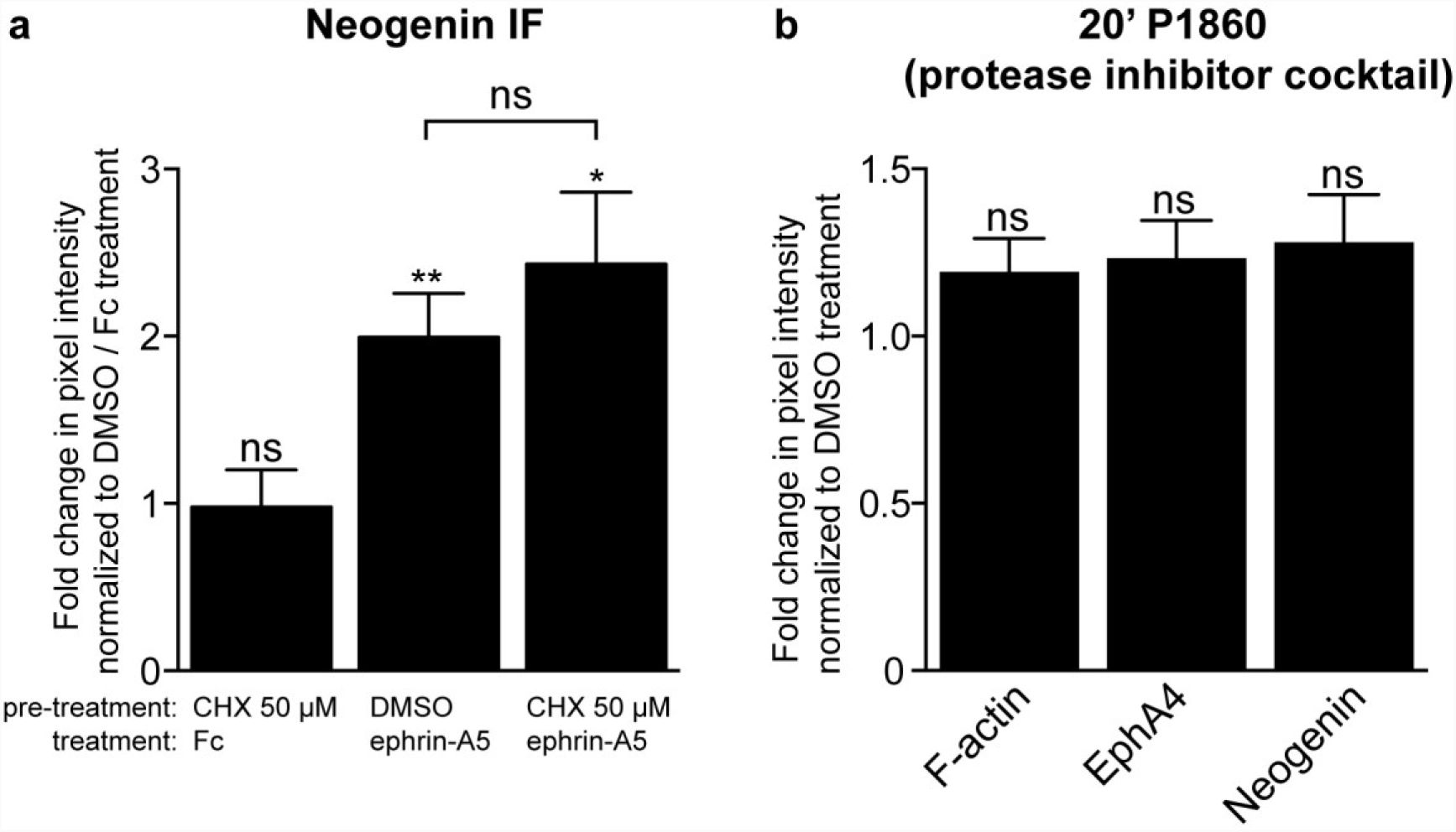
The ephrin-A5 induced increase in Neogenin protein levels in LMC growth cones may occur independently of protein synthesis and proteolytic degradation. (a) Quantification of Neogenin IF in LMC growth cones pre-treated for 20’ with either DMSO or CHX 50 µM followed by a 5’ treatment with either Fc or ephrin-A5 500 ng/mL. The inclusion of CHX did not significantly change Neogenin IF in growth cones treated with ephrin-A5 (p=0,4177). (b) Relative to DMSO, a 20’ treatment with a protease inhibitor cocktail (P1860, 1/200) failed to increase the levels of F-actin (p= 0.0828), EphA4 (p= 0.0647) and Neogenin IF (p= 0.0769) in growth cones of LMC explant cultures. Data are shown as mean ±SEM, statistical significance was tested using a two-tailed unpaired sample t-test. (a,b) N=4.

### The cytoplasmic tail of EphA4 is dispensable for potentiating the ephrin-A5-induced increase in Neogenin abundance

Next, we asked if the intracellular domain of EphA4, required for the relay of signals in ephrin:Eph forward signalling^52^, is also required for the EphA4-induced potentiation of ephrin-A5-mediated Neogenin IF increase. To do so, we explanted LMC neurons from chick spinal cords electroporated with either expression plasmids encoding an EphA4 and GFP fusion protein (*EphA4-GFP*) or a truncated EphA4 missing the intracellular domain and GFP fusion protein (*EphA4ΔICD-GFP*)^10^. EphA4 immunofluorescence and GFP fluorescence was used to confirm fusion protein expression levels (**Figure 8b,c**). Both sets of explants were treated with either MN media or ephrin-A5, and Neogenin IF levels were quantified in growth cones (**Figure 8a**). In line with ephrin-A5-led induction of EphA4 receptor clustering, in EphA4-GFP-expressing growth cones, ephrin-A5 exposure led to the formation of patches of GFP signal, that also co-localized with Neogenin (**Figure 8e-g**). In contrast, in growth cones expressing EphA4ΔICD-GFP and treated with ephrin-A5, GFP signal was more diffuse, in line with the requirement for the intracellular domain of EphA4 for ephrin-A5 induced EphA4 clustering (**Figure 8e,k**)^30^. Surprisingly, EphA4ΔICD-GFP-expressing growth cones exposed to ephrin-A5 showed a robust increase in Neogenin IF when compared to controls (**Figure 8a,j,m**, 6.6-fold induction, p=0.0007), which was indistinguishable from that observed in EphA4-GFP expressing growth cones (**Figure 8a**,**d,g**, 5.1-fold induction; p=0.3). These results suggest that the ephrin-A5 increase in Neogenin IF does not require canonical EphA signalling through the EphA4 cytoplasmic tail.

**Figure 8.**
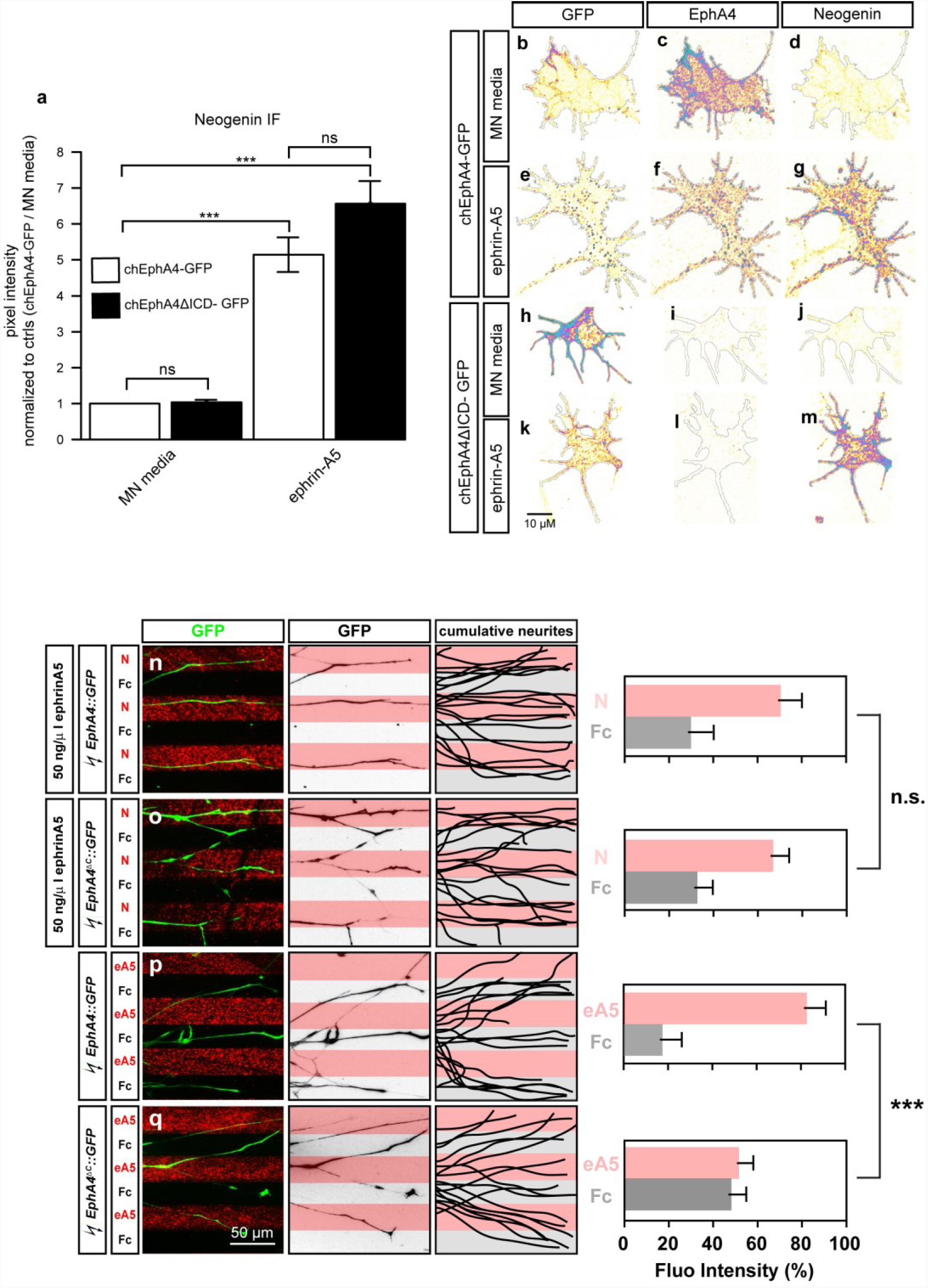
The cytoplasmic tail of EphA4 is dispensable in potentiating the ephrin-A5 induced increase in Neogenin protein levels and sensitization of LMC axons to netrin-1. (a-m) LMC explants from chick spinal cords electroporated with either a chEphA4-GFP or chEphA4ΔICD-GFP expression plasmids were subject to a 20’ treatment of either MN media or ephrin-A5 at 100 ng/mL and immunostained for Neogenin (a) Quantification of Neogenin IF in growth cones shows that the ephrin-A5 induced increase in Neogenin signal occurs in growth cones expressing both plasmids (chEphA4-GFP: 5.14-fold increase, p=0.0010, N=3; chEphA4ΔICD-GFP: 6.56-fold increase p=0.0035, N=4). (b-m) Examples of GFP, EphA4 and Neogenin IF in growth cones quantified in (a). (n-q) Growth preference on protein stripes exhibited by LMC axons. Left panels: explanted LMC neurites expressing chEphA4-GFP and chEphA4ΔICD-GFP on netrin-1 (N)/Fc stripes bath treatment of ephrin-A5 (n,o) or ephrin-A5 (eA5)/Fc stripes (p,q). Middle panels: inverted images of GFP signals shown at left panels. Right panels: superimposed images of five explants from each experimental group representing the distribution of GFP^+^ LMC neurites. Quantification of lateral LMC neurites on first (pink) and second (pale) stripes expressed as a percentage of total GFP signals. Noted that both chEphA4-GFP and chEphA4ΔICD-GFP expressed LMC neurites show preferences over Netrin 1 stripes. Minimal number of neurites: 85. Minimal number of explants: 13. N, netrin-1; eA5: ephrin-A5; error bars = SD; *** = P<0.001; statistical significance computed using Mann-Whitney U test.

### The cytoplasmic tail of EphA4 is dispensable for the sensitization of LMC axons to netrin-1 by ephrin-A5

Finally, to test whether the cytoplasmic tail of EphA4 is required for the functional ephrin-A5-induced sensitisation of LMC axons to netrin-1, we challenged the above LMC neurons expressing EphA4-GFP or EphA4ΔICD-GFP with ephrin-A5/Fc stripes (**Figure 8p,q**). While LMC axons expressing EphA4-GFP showed robust avoidance of ephrin-A5 stripes, this effect was diminished in LMC axons expressing EphA4ΔICD-GFP, in line with the requirement of the intracellular tail of EphA4 for forward ephrin-A:EphA signalling (**Figure 8p,q;** EphA4-GFP: ephrin-A5 stripe = 17%, EphA4ΔICD-GFP: ephrin-A5 stripe = 52%; p<0.001) Intriguingly, the expression of EphA4-GFP and EphA4ΔICD-GFP resulted in the same enhancement of growth preference over netrin-1 stripes in the presence of bath-applied ephrin-A5 (**Figure 8n,o**; EphA4-GFP: netrin-1 70 ± 10%, EphA4ΔICD-GFP: netrin-1 67 ± 7%; p=0.2). Together, these data argue that EphA4 can promote ephrin-A5-mediated sensitization of LMC axons to netrin-1 in the absence of its cytoplasmic tail.

## DISCUSSION

Our experiments provide evidence that ephrin-A5:EphA4 interaction sensitizes spinal motor axons growth cones to netrin-1 by increasing the abundance of Neogenin, thus enhancing their netrin-1 binding and guidance responses. Further evidence suggests that these effects occur outside of canonical ephrinA:EphA signalling. Below, we discuss the potential molecular mechanisms underlying these findings, and, their *in vivo* relevance to motor axon guidance, and other functions of Neogenin and netrin-1 signalling.

### Mechanisms of ephrin-A5 induced Neogenin upregulation

While our experiments strongly support the idea that ephrin-A5 exposure of LMC growth cones results in increased surface levels of Neogenin, we have yet to develop a complete mechanistic understanding of this phenomenon. The time-frame of the effect is on the order of minutes, and compatible with the idea of an axon guidance cue causing a rapid increase in growth cone protein translation^53,54^. However, the ephrin-A5-evoked increase in growth cone Neogenin occurs in the presence of protein synthesis inhibitors, arguing against the involvement of protein synthesis. Suppression of Neogenin degradation also does not appear to be involved since a variety of protease, proteasome and lysosomal degradation inhibitors failed to increase Neogenin levels. One remaining possibility is that cell surface Neogenin levels are controlled through its trafficking. Our data exclude the cAMP and PKA – dependent exocytosis that controls DCC levels at the cell surface^55^. In Cos-7 cells overexpressing a fluorescently-tagged EphA2, ephrin-A1 induces a rapid translocation of EphA2 from Rab11+ recycling endosome to the plasma membrane, followed by a reduction in EphA2 cell membrane levels and its appearance in Rab5+ late endosomes^29^. Considering that this experiment was performed using ephrin-A1 at 2 µg/mL whereas our experiments were mostly done at an ephrin-A concentration 20 times lower, it is possible that treatment with lower concentrations of ephrin-As would result in a prolonged increase in plasma membrane levels of EphA4 and less robust endocytosis^56^. In this context, we can hypothesize that in LMC growth cones, ephrin-A5 could induce the trafficking of EphA4-carrying vesicles to the growth cone cell membrane, and if such vesicles also contain Neogenin, this might result in increased cell surface Neogenin levels.

Our results show that the ephrin-A5-induced Neogenin upregulation is potentiated by EphA4 and depends on its ephrin ligand binding domain but, surprisingly, not its intracellular domain. Ephrin-A5 increased the co-localisation of Neogenin and EphA4 in LMC growth cones suggesting that the elevation of surface Neogenin may be local and occurring preferentially in the vicinity of ephrinA5:EphA4 interactions. The EphA4 intracellular domain contains a signalling-essential tyrosine kinase domain^52^ and is essential for ephrin-A5-induced EphA4 clustering^30,57^. EphA4 lacking this domain did not form clusters when exposed to ephrin-A5, but its expression was sufficient to elevate Neogenin levels in LMC growth cones suggesting that the EphA4 intracellular domain and clustering are dispensable for Neogenin elevation. Furthermore, since the cytoplasmic tail of Eph receptors is also required for their efficient signalling through endocytosis^58^ it is likely that the ephrin-A5:EphA4 signalling events leading to increased Neogenin, are initiated at the plasma membrane and are distinct from a canonical Eph signalling cascade. One possibility is that EphA4 participates in a molecular complex that includes a receptor that can transfer a signal initiated by ephrin-A5-EphA4 binding through its own cytoplasmic tail. An example of this, is the reverse Eph:ephrin signalling that involves signal transfer to the c-Ret receptor, found in a complex with ephrin-A5 and required for normal LMC axon guidance^12,13^.

Despite their relatively high levels of EphA4, dorsal spinal cord growth cones did not respond to ephrin-A5 by increasing Neogenin levels, suggesting that EphA4 is not sufficient for this effect. Ephrin-A5 also failed to increase EphA4 levels in dorsal spinal cord growth cones suggesting that the molecular machinery behind the ephrinA5:EphA4 increase in EphA4 present in LMC neurons may be absent in dorsal spinal cord explants, possibly due to the much higher basal levels of EphA4 in growth cones of dorsal explants *in vitro*. It is thus feasible that the increase in Neogenin seen in LMC growth cones may be molecularly linked to the increase in EphA4 levels. This idea is in line with the hypothesis of ephrin-A5 inducing the trafficking of EphA4+/Neogenin+ vesicles towards the cell membrane in LMC growth cones. Combinatorially-expressed growth cone guidance signal receptors can modulate netrin-1 axon guidance. For instance, most thalamocortical axons do not normally respond to netrin-1, however, a subset of them can be attracted towards netrin-1, if it is provided in combination with the Robo ligand Slit1^23,59^. This effect depends on PKA, and the expression of Robo1 and its co-receptor FLRT3 whose signalling causes an increase in the levels of DCC on the growth cone surface^24^. A similar requirement for a combination of receptors may apply to the signals that result in ephrin-A5-mediate induction of Neogenin levels. A biochemical fraction of embryonic brain cell membranes, termed Netrin-synergising activity (NSA), containing a protein (or proteins) of a molecular weight of 25-35 kDa, synergises with netrin-1 to induce axonal outgrowth from dorsal spinal cord explants^60^. NSA and ephrin-A5 share a predicted molecular weight of ∼26 kDa, but ephrin-A5’s predicted isoelectric point of 5.97, which does not align with the prediction that the NSA is a basic protein^61^.

### Ephrin-A5-induced Neogenin upregulation in motor axon trajectory selection

The choice of limb trajectory made *in vivo* by LMC neurons is also consistent with the induction of Neogenin by ephrin-A5. LMC motor neurons express Neogenin and DCC attractive netrin receptors (NetrinR^attractive^) and can be divided into two populations according to their EphA4 expression. Lateral LMC neurons express high levels of EphA4, are attracted to netrin-1 *in vitro*, and grow towards netrin-1-expressing dorsal limb mesenchyme, away from ephrin-A5-expressing ventral limb mesenchyme. Based on our findings, it is plausible that in addition to repelling EphA4-expressing lateral LMC axons into the dorsal limb, ephrin-A5 in the ventral limb induces a high level of Neogenin expression in lateral LMC axons that stray there by mistake, making them more responsive to netrin-1, and allowing them to choose the correct limb trajectory even in mice with *Neogenin* or *netrin1* hypomorphic mutations^18^. In this context, it is worth noting that our experiments also demonstrate a Neogenin induction by ephrin-B2, which is normally present in the dorsal limb and is consistent with lateral LMC axons targeting this limb domain.

Medial LMC neurons, have negligible EphA4 expression and are repelled from netrin-1 through its receptor Unc5c, resulting the choice of a ventral limb trajectory ^18^. Expression of EphA4 in medial LMC neurons is sufficient to redirect them towards the dorsal limb and has been explained as resulting from EphA4-mediated repulsion from ventral limb ephrin-A5. This effect could be potentiated by ephrin-A5-induced upregulation of Neogenin in medial LMC neurons, resulting in their attraction to dorsal limb netrin-1. Medial LMC neurons also express Unc5c which can mediate repulsion from netrin-1 by itself or in conjunction with NetrinR^attractive^ receptors^14^. Elevation of Neogenin levels could result in increased abundance of NetrinR^attractive^ – only complexes and fewer Unc5c/ NetrinR^attractive^ complexes, leading to greater attraction towards dorsal limb netrin-1.

### Potential developmental functions of ephrin-A5-induced Neogenin upregulation

Loss of Neogenin, the Neogenin ligands RGMa, RGMb, and netrin-1 cause neural tube closure defects and have been proposed to be due to decreased cell-cell adhesion at the dorsal aspect of this structure^62-64^. Similar defects are also present in EphA7 and ephrin-A5 null embryos^65^ and were hypothesised to reflect the adhesive function of an EphA7 splice isoform that lacks the intracellular domain^65^. However, in light of our results, ephrin-A5-EphA7 interactions in the neural folds could be contributing to neural tube closure by increasing Neogenin cell surface levels and thus promoting Neogenin-mediated adhesive interactions.

Post-synaptic EphA4 and ephrin-A3 expressed by astrocytes are required for synaptic plasticity by modulating long term potentiation (LTP) in the hippocampus^66^. This requirement for EphA4 for LTP occurs independently from its cytoplasmic tail, since LTP deficits seen in EphA4 null mice are absent in mice expressing EphA4 lacking its cytoplasmic tail^67^. Loss of EphA4 increases the abundance of glial glutamate transporters and LTP deficits can be rescued by blocking glial glutamate transporters^66^. The mechanism underlining the requirement for EphA4 in LTP is unknown^66^. Interestingly, *Dcc* null mice also show LTP deficits in the hippocampus, proposed to originate from decreased levels of Dcc-dependent Src activation of NMDA receptors^68^. The overlapping functions between chicken Neogenin and mouse Dcc as well as the expression of Neogenin in the hippocampus in mice raises the possibility that ephrin:EphA4 interactions may in part promote LTP by increasing the abundance of post-synaptic Dcc/Neogenin. The Ephrin-A induced increase in post-synaptic Dcc/Neogenin could result in higher levels of Src dependent NMDA receptor phosphorylation and LTP induction^18,32,69^.

### Conclusion

Together, our data reveal a novel interaction between classical axon guidance ligands and their receptors. Ephrin-A5-directed increase of Neogenin levels in motor neuron growth cones does require EphA4 but does not appear to proceed through a canonical Eph signalling mechanism. It also results in augmented axon guidance responses to netrin-1 that are consistent with the genetically-tested requirements for ephrin and netrin signalling in motor axon guidance *in vivo*. Given the importance of netrin and Neogenin signalling inside and outside the nervous system, this new paradigm could have implications in a wide variety of biological processes.

## Materials and methods

### Explant culture

All animal experiments were carried out in accordance with the Canadian Council on Animal Care guidelines and approved by the IRCM Animal Care Committee and the McGill University Animal Care Committee. Fertilised chicken eggs (FERME GMS, Saint-Liboire, QC, Canada) were incubated (Lyon Technologies, model PRFWD) at 39°C according to standard protocols^25^. LMC explants were collected from HH st. 24-26 lumbar spinal cords and incubated in 95% air and 5% CO2 at 37°C in MN medium for about 18 hours as previously described^27^. 20 mL of MN medium solution is: 19.2 mL of Neurobasal medium (Invitrogen, cat. no. 21103-049), 400 µL Serum-free supplement (50×, B-27; Invitrogen, cat. no. 17504-044), 2 µL of l-glutamic acid (50 mM, Sigma-Aldrich, cat. no. G8415), 73 mg of l-glutamine (Invitrogen, cat. no. 21051-024) and 200 µL of penicillin-streptomycin (100×, Invitrogen, cat. no. 15140-122). Prior to explant culture, tissue culture dishes (Sarstedt, cat. no. 83.3901.300) were coated with 20 µg/mL Laminin (Invitrogen, cat. no. 23017-015) for 2 hours at 37°C and rinsed with Neurobasal medium.

### Explant treatment reagents

For inhibitor treatments, half of the motor neuron media in the explant cultures was replaced with media containing the inhibitor or dimethyl sulfoxide (DMSO; Sigma-Aldrich, cat. no. D2650) and incubated for 20’ prior to cue treatment. The following drugs were used: cycloheximide, P1860 protease inhibitor cocktail, KT5720, KT5823, γ-Secretase Inhibitor IX (DAPT), Anisomycin and SU6656 were purchased from Millipore Sigma. MG132, Chloroquine diphosphate, RVKR (Deconoyl-RVKR-CMK) and the KYL peptide were purchased from Tocris.

For ephrin and netrin treatments, half of the culture media was replaced with media alone or media containing recombinant mouse netrin-1 (R&D systems cat. no.1109-N1/CF), recombinant human ephrin-A5-Fc (R&D systems cat. no. 374-EA), recombinant mouse ephrin-B2-Fc (R&D systems cat. no. 496-EB), recombinant mouse ephrin-A2-Fc (R&D systems cat. no. 8415-A2) or Fc (Millipore Sigma cat. no. 401104). Prior to explant treatment, recombinant ephrins or Fc were preclustered in a 5:1 molar ratio with either mouse or goat anti-human Fc (Millipore Sigma cat. no. 16760 and 12136 respectively) for 30’ at 37°C.

### Chick *in ovo* electroporation

Chicken *in ovo* electroporations were carried out as previously described^34^ at HH st.18–19 and harvested at HH st. 24-25. Chicken embryos were electroporated with either the *pN2-eGFP* (Invitrogen) expression plasmid alone or in a 1:4 molar ratio combination with *pCAGGS-mEphA4* ^70^ or with *pN2-chEphA4-GFP or pN1- chEphA4ΔICD- GFP* (cytoplasmic domain deleted) plasmids^10^.

### *In vitro* stripe assays

In vitro stripe assay using explants of spinal motor columns were performed as described^27^. In brief, carpets of alternating stripes of Netrin-1, ephrin-A5-Fc, or Fc only as controls were prepared using silicon matrices with micro-well system (provided by Dr. Martin Bastmeyer’s laboratory). E5 chick spinal motor column was dissected using sharp tungsten needles (World Precision Instruments) and collected in MN medium. The excised motor column was then trimmed into explants with the size of 1/4 width of motor column, and 20 explants were plated on laminin coated culture dishes containing different combinations of stripe carpets in motor neuron medium and incubated overnight. Following incubation, motor column explants were fixed with 1:1 mixture of 4% paraformaldehyde (Sigma) and 30% sucrose in PBS for 5 minutes followed by 4% paraformaldehyde for 5 minutes. After PBS washes, explants were incubated with selected primary antibodies diluted in blocking solution containing 20% serum in 0.3% Triton-X/PBS(Sigma) for 2 hrs at room temperature (RT). Following PBS washes, explants were incubated with secondary antibodies diluted in the same blocking solution for 2 hrs at RT.

### Immunohistochemistry

Prior to immunostaining, explants were fixed by replacing half of the culture media with a 37°C solution of 4% PFA, 3% sucrose in PBS for 20’ at RT and washed repeatedly with PBS. Primary antibodies were incubated in blocking solution (1% heat-inactivated horse serum in 0.1% Triton-X/PBS; Millipore Sigma) either 1 hour at 37°C or overnight at 4°C. The following primary antibodies were used: goat anti-Neogenin (1:300; R&D cat. no. AF1070), rabbit anti-Neogenin (1:2000; Abcam cat. no. 190263), rabbit anti-EphA4 (1:1000; Santa Cruz Biotechnology cat. no. S20), rabbit anti-Netrin1 (1:1000; Abcam cat. no. EPR5428), guinea-pig anti-Unc5c (1:500; Thomas Jessell lab.), mouse anti-Neuronal Class III β-Tubulin (Tuj1) (1:2000; Covance cat. no. MMS-435P), mouse anti-chicken BEN (1:100; Developmental Studies Hybridoma Bank). For surfaced enriched staining, antibodies were added simultaneously with the cue treatment solution. Explants were washed repeatedly with PBS prior to incubation with appropriate secondary antibodies in blocking solution for 1h at RT. The following secondary antibodies were used: Cy3- (or Cy-5)-conjugated AffiniPure donkey anti-mouse (rabbit, goat, or guinea pig) IgG (1:1000 for Cy3, 1:500 for Cy5 secondary antibodies; Jackson ImmunoResearch Laboratory), Alexa Fluor 488 donkey anti-mouse (rabbit or sheep) IgG (1:1000; Invitrogen). F-actin was detected using Alexa Fluor™ 568 Phalloidin (1:300; Thermo Fisher scientific). Explants were then washed repeatedly with PBS and mounted with Mowiol mounting medium (9.6%, w/v Mowiol (Calbiochem), 9.6% (v/v) 1 M Tris–HCl (Fisher Scientific), 19.2% (v/v) Glycerol (Fisher Scientific), in H2O).

### Image acquisition and quantification

16-bit tiff images were acquired using LSM700 and LSM710 Zeiss confocal microscopes. Regions of interests (ROIs) delimitating growth cones and subsequent analysis was done using ImageJ. The ROI areas and mean pixel intensity within the ROIs was determined using the measurement tool in ImageJ. Images of growth cones depicted in figures were subject to a rainbow RGB lookup table in ImageJ followed by the overlay of a difference mask in Adobe illustrator. To assess fluorescence distribution within the growth cones (**Figure 4g**), a MATLAB application as designed and programmed by Dr. Dominic Fillion (IRCM). The application divides growth cones into 100 bins, with bin 1 at the geometric centre and bin 100 following the outer perimeter of the growth cone. The mean fluorescence value for each bin was determined and plotted, bin#100 was omitted from the analysis due to the region of interest (ROI) occasionally slightly over representing the growth cone resulting in the absence of signal in bin#100. To analyse fluorescence overlap (**Figure 6a,m**), the fluorescence signal in individual channels was thresholded and an image representing signal overlap between two channels was generated and the mean pixel intensity was quantified. To determine the change in overlap, the data was treated as such ((% signal overlap with cue treatment) / (%signal overlap control treatment).

### Statistical analysis

Data from the experimental replicate sets were evaluated using Microsoft Excel and Aabel (Gigawiz), statistical significance was set at 0.05. Standard error of the mean values, p values and the number growth cones analysed can be found in the Supplementary Excel file.

## Author contributions

Experiments were designed by L.-P. Croteau under the supervision and input from A. Kania. The stripe assay experiments were performed and analysed by T.- J. Kao. All other experiments were performed and analysed by L.-P. Croteau. The manuscript was written by L.-P. Croteau and A. Kania.

### Acknowledgments

We would like to thank E. Pasquale for providing us with the *EphA4ΔICD-GFP* and *EphA4-GFP* expression plasmids. We also thank D. Fillion for programming the MATLAB application used to analyse the immunofluorescence distribution within growth cones and C. Law for programming the ImageJ macro used to define the region of interest delimiting growth cones. We also thank M. Liang for performing the chick *in ovo* electroporations. L.-P. Croteau was supported by a doctoral training award from the Fonds de recherché du Québec – Santé (26337). This work was supported by grants to A. Kania from the Canadian Institutes of Health Research (MOP-97758, MOP-77556), Brain Canada, Canadian Foundation for Innovation, and the W. Garfield Weston Foundation. A. Kania was also supported by an FRSQ Chercheur-boursier Senior career award.

## Additional Information

## Competing interests

The authors declare no financial or non-financial competing interests.

## Data Availability

The MATLAB application used for the analysis of IF distribution within growth cones as well as the ImageJ macro used to create the ROIs may be provided upon request.

---

**Supplementary figure 1.**
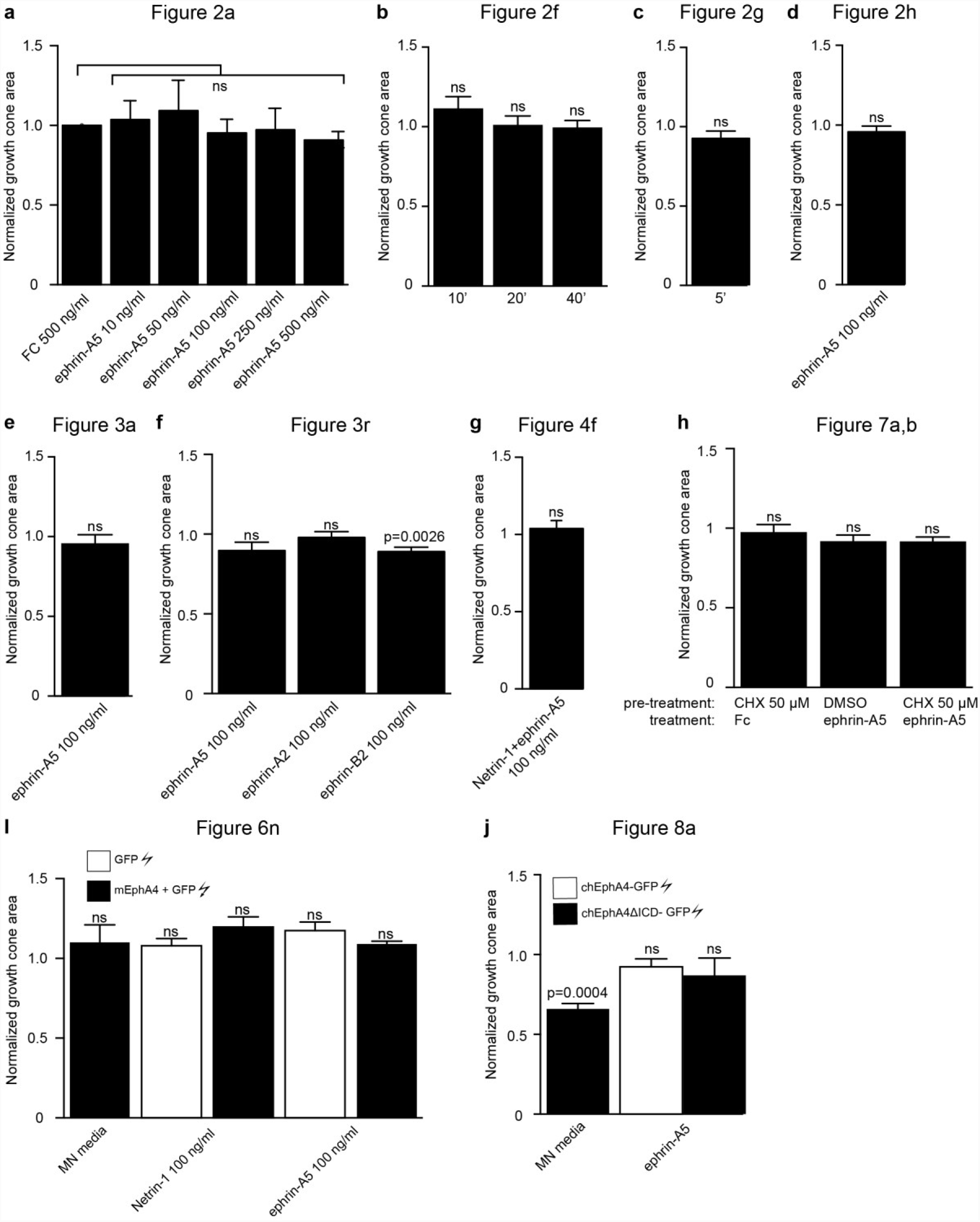
Mean area measurements of growth cones selected for IF analysis. (f) Compared to a 20’ Fc 100 ng/mL treatment, ephrin-B2 100 ng/mL results in a 0.89-fold decrease in mean growth cone area (p=0.0026). (j) Compared to growth cones overexpressing EphA4-GFP, overexpression of EphA4ΔICD-GFP resulted in a 0.66-fold decrease in mean growth cone area when treated with MN media for 20’ (p=0.004). In all other instances, mean growth cone measurements did not differ from controls. (p>0.05). Data are shown as mean ±SEM, statistical significance was tested using a two-tailed unpaired sample t-test.

**Supplementary figure 2.**
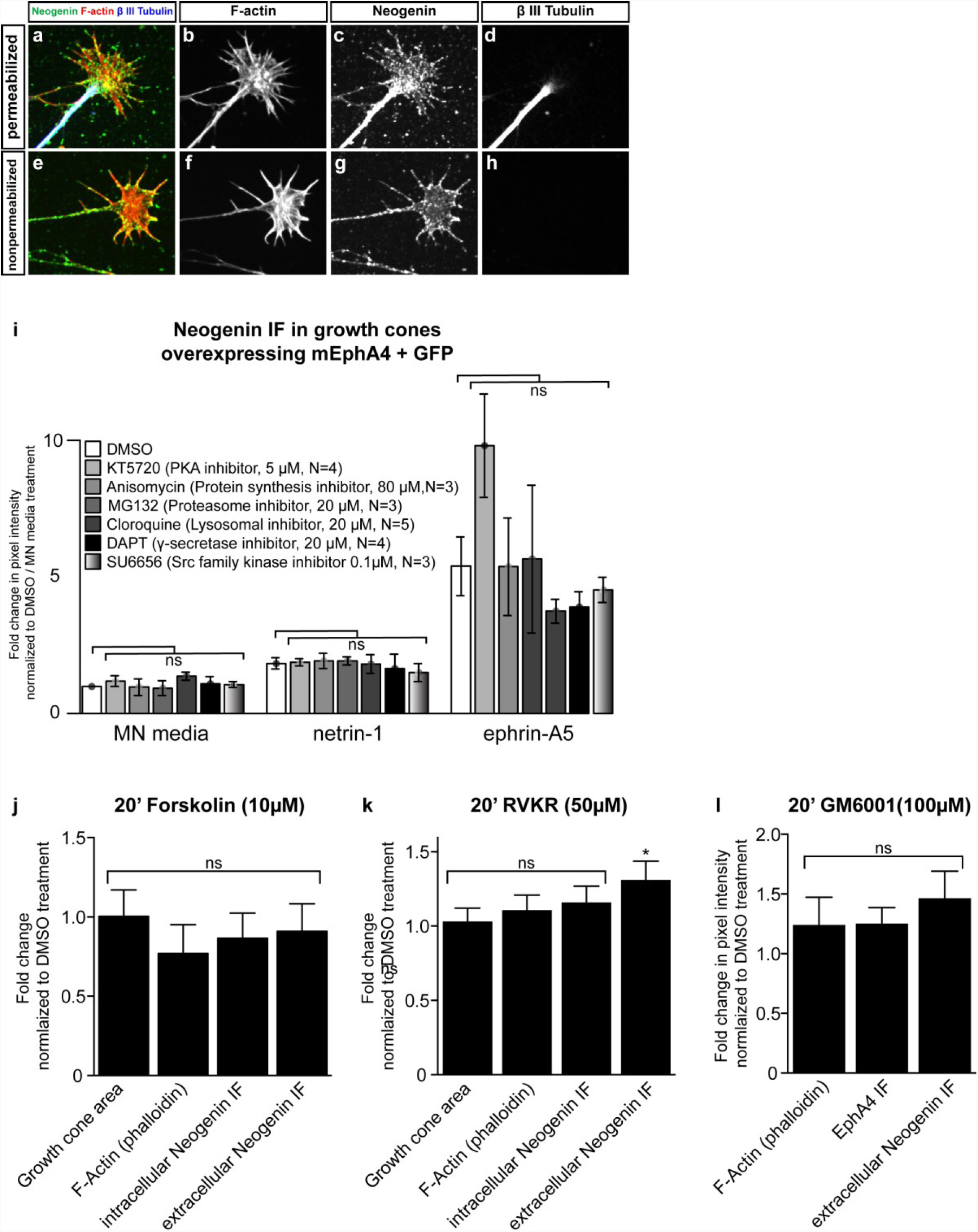
Surface enrichment of Neogenin IF and the effects of various drugs on Neogenin IF in LMC growth cones. (a-h) To confirm the surface enrichment of Neogenin when polyclonal anti-Neogenin antibody is added to live cultures (Figure 4a-c), an antibody against the intracellular protein βIII tubulin was included in fixed (a-d) and live (e-h) cultures, secondary antibodies and phalloidin to detect F-actin were added post-fixation (a-h). (i) The ephrin-A5 induced increase in Neogenin IF prevails despite inhibiting specific cell functions. LMC explants from chick spinal cords electroporated with a GFP expression plasmid in combination with a mEphA4 expression plasmid were incubated in the presence of various drugs for 20’ prior to a 20’ incubation with either MN media, netrin-1 or ephrin-A5 at 100 ng/mL followed by immunostaining for Neogenin. Drug treatments failed to modify Neogenin IF in GFP+ growth cones. (j-l) LMC explant cultures were treated for 20’ with either the adenylate cyclase activator Forskolin (j), the proprotein convertase inhibitor RVKR (k) or the metalloprotease inhibitor GM6001(l) followed by a quantification of F-actin, EphA4 and Neogenin IF in LMC growth cones. Relative to DMSO treatment, RVKR treatment results in 1.31-fold increase Neogenin IF (p=0.034), all other measurements did not differ significantly (p>0.05). Data are shown as mean ±SEM, statistical significance was tested using a two-tailed unpaired sample t-test. (j) N=4, (k) N=6, (l) N=5.

